# A pan-ontology view of machine-derived knowledge representations and feedback mechanisms for curation

**DOI:** 10.1101/2021.03.02.433532

**Authors:** Tomasz Konopka, Damian Smedley

## Abstract

Biomedical ontologies are established tools that organize knowledge in specialized research areas. They can also be used to train machine-learning models. However, it is unclear to what extent representations of ontology concepts learned by machine-learning models capture the relationships intended by ontology curators. It is also unclear whether the representations can provide insights to improve the curation process. Here, we investigate ontologies from across the spectrum of biological research and assess the concordance of formal ontology hierarchies with representations based on plain-text definitions. By comparing the internal properties of each ontology, we describe general patterns across the pan-ontology landscape and pinpoint areas with discrepancies in individual domains. We suggest specific mechanisms through which machine-learning approaches can lead to clarifications of ontology definitions. Synchronizing patterns in machine-derived representations with those intended by the ontology curators will likely streamline the use of ontologies in downstream applications.

## Introduction

Ontologies are schemes that conceptualize, systematize, and organize knowledge. In biomedicine, the term “ontology” has also been used in a practical sense to refer to a collection of definitions relevant to a research domain and the relationships between them^1,2^. A prominent example is the Gene Ontology^3^ that organizes gene functions, but hundreds of other ontologies provide infrastructure to coordinate databases and power bioinformatic applications^1^. In parallel to the adoption of ontologies, advances in machine-learning (ML) have demonstrated the possibilities of creating representations of knowledge from labeled datasets or even from raw data^4–6^. The overlap between curated structures and automated pattern discovery raises the need to clarify the relation between those components. Indeed, because they serve distinct purposes, it is of interest to elucidate how ontologies can enhance ML models, and, conversely, in what way should insights from ML influence the development of ontologies. Ontologies are used across the spectrum of biomedical research, so it is also relevant to understand how observations about ontologies and ML in one domain can be transferred to others.

With regard to the overlap between ontological- and machine-driven approaches, there is a compelling argument that they are complementary rather than competing alternatives. This stems from the need to integrate information from distinct sources into a common pool of knowledge. Repositories such as the OBO foundry^7^, the Ontology Lookup Service^8,9^, and the BioPortal^10,11^ host ontologies that span research domains from chemistry, through molecular biology and model organisms, to ecology. These projects provide four distinct components: vocabularies of terms that encompass their research domains, text definitions for those terms, stable identifiers, and hierarchical relationships between the terms^1^. Those components are transparent to human researchers, and have been proven to be adaptable and extendable. They provide a framework for integrating new datasets with existing databases. Because the definitions can also be processed by machine agents, ML applications can use ontologies to extract data from databases, use labeled datasets to train models, and report outputs in an interpretable fashion. Ontologies thus enable ML models to participate in knowledge-building with human curators as well as with other machine-driven approaches.

The topic of how ontologies can enhance machine learning applications is a rich one, especially because implementations can utilize one or more of the four distinct information components (vocabularies, definitions, identifiers, and relations) as well as external associations (e.g. between genes and ontology identifiers). The text components of ontologies have long been used for text mining^12^. Vocabularies and identifiers are used as labels for ML classifiers, especially in applications where the labels are numerous and need to be managed carefully. A recent example in this domain used neural networks to annotate text with ontology-based subject labels^13^. Another active direction in machine learning research relevant to ontologies has been representation learning. An influential technique in this area has been the encoding of words into an abstract low-dimensional space^14^. Analogous encodings for ontology terms^15–18^, also referred to as embeddings, have been evaluated in the context of the gene ontology^19–21^. The embeddings transform concepts into a small number of coordinates, thereby simplifying the training of downstream models. For example, embeddings of ontology-based features in ML classifiers can improve predictions of protein-protein interactions.

In the reverse direction, i.e. regarding the use of computational tools to enhance ontologies, research activity has included auditing^1,22,23^. Approaches have exploited regularities in the naming conventions in SNOMED^24^ and GO to suggest missing links between existing ontology terms^25–28^. These case-studies demonstrate that the curation of ontologies may benefit from automated tools, but their rule-based approaches cannot be easily generalized. Furthermore, the impact of knowledge representations built by machine learning techniques on the development of ontologies is an unexplored area. It is unclear, for example, whether neural networks trained on biomedical corpora^13,29^ can feed insights back into the definitions of ontology-based labels. Similarly, it is unclear whether embeddings of ontology terms into low-dimensional spaces, while useful for downstream applications, can help guide development of ontology structures. These issues are arguably open because ML studies have regarded ontologies as static annotations rather than evolving structures that can adapt to correct any flaws. Indeed, the transfer of knowledge from machine-derived representations into ontology definitions may be an avenue toward streamlining the maintenance of ontologies and generating long-term impact from ML studies.

Ontologies are used in several research domains and the interaction between curation and ML is relevant to all of them. Past analyses of the pan-ontology landscape have reviewed structural aspects ontologies, but without direct relation to ML. For example, the extent of overlap between ontology projects^30,31^ has been quantified, and has been shown to impact how researchers can find ontology terms to annotate a dataset^32^. From a different perspective, the properties of ontology graphs have been shown to vary substantially across projects^33^. The graph structure is utilized during training phases of ML models, for example to construct node embeddings, so this variability has the potential to affect downstream applications. The effects have been exemplified through case studies with the Gene Ontology^19–21^, but it is unclear to what extent observations can be generalized to other ontologies and to other domains. Text annotations associated with ontology terms can also be used for ML training^17^ and are crucial to the interpretability of ontology terms, but these too have not been systematically assessed.

Here, we performed a review of open biomedical ontologies to elucidate interconnections between human-curated ontologies and their ML-derived representations. Part of our work aimed to extend explorations of representation learning to a diverse gamut of ontologies. We placed emphasis on the idea that ontologies are not static sources of ground-truth annotations. This viewpoint allowed us to employ ML to assess internal consistency within individual ontologies, in particular between text-based definitions and ontology hierarchies. We also explored mechanisms through which machine-driven representations can translate into changes in ontology structures.

## Results

### Biomedical ontologies from diverse domains share common traits

Aiming to cover the spectrum of biomedical research domains, we compiled a list of ontologies by polling the Open Biomedical Ontologies Foundry^7^ and the Ontology Lookup Service^8^. We downloaded source files and converted them into a single common format (Methods). This resulted in a set of 194 ontologies. We then extracted metadata, term identifiers, term names, term definitions, and ‘is a’ relationships that organize terms (also called classes) into the hierarchical structures often visualized in ontology browsers. These features are common to all ontologies and can thus serve as the basis of a pan-ontology analysis.

The sizes of the ontologies spanned six orders of magnitude (Figure 1A), but the majority consisted of between 100 and 10,000 terms. The time intervals between our downloads and the last file updates were also diverse (Figure 1B). 45 ontologies were not updated for more than two years. This may indicate either a lack of active maintenance, or that those ontologies already cover their intended use cases^34^. For 42 ontologies, the time interval could not be determined from the downloaded data. Separately, we summarized the ontology graph structure by counting the number of hierarchy roots defined by ‘is a’ relationships (Figure 1C). Only 39 ontologies adhered to a simple hierarchical structure with a single root node. The majority of projects organized terms under two or more roots, and 57 contained more than 10 roots. This complexity may be a result of an intentional separation of data into disjoint groups (e.g. into ‘biological process’, ‘molecular function’, and ‘cellular component’ in GO), but can also arise when ontologies reuse components from other projects.

**Figure 1.**
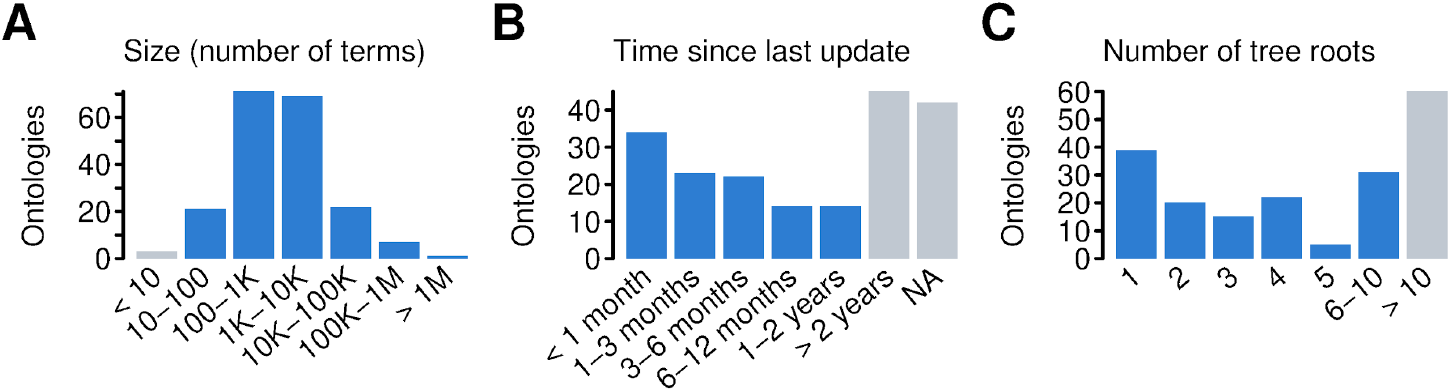
Summary of biological ontologies. Panels show a collection of ontologies stratified by (A) the number of terms (classes), (B) the time since last update, and (C) the number of roots in the hierarchy. Ontologies in light-colored strata were excluded from subsequent analysis.

Using the above global properties as filtering criteria, we identified a group of 75 ontologies that had at least 10 terms, had been updated within the last two years, and contained fewer than 10 root nodes. We reasoned that this selection would place emphasis on projects that are under active development and therefore most likely to incorporate any insights based on machine learning. By focusing on hierarchies with a small number of root nodes, we attempted to reduce confounding effects due to disjoint or complex graphs.

After selecting the group of 75 ontologies for in-depth analysis, we characterized their internal features (Figure 2, Supplementary Figures S1, S2). Several metrics about the structural aspects of ontologies have already been reviewed^33^, so we only investigated a small number of features. With more than two million terms, the largest ontology was *NCBITAXON -* a taxonomy of organisms and species^35^. This was an outlier and a ranking of other ontologies displayed a smooth gradation in terms of size. We also computed the distribution of term depths normalized by the logarithm of the size of each ontology. Noting that a balanced binary tree produces a distribution centered around 0.8, this revealed that most ontologies are much shallower than such a reference tree. Furthermore, distributions of the number of parents indicated that most ontologies include terms with more than one asserted parent. Indeed, in 22 of the 75 selected ontologies, at least 25% of terms had more than one parent. Thus, these structures routinely encode concepts that bridge distinct ontological branches.

**Figure 2.**
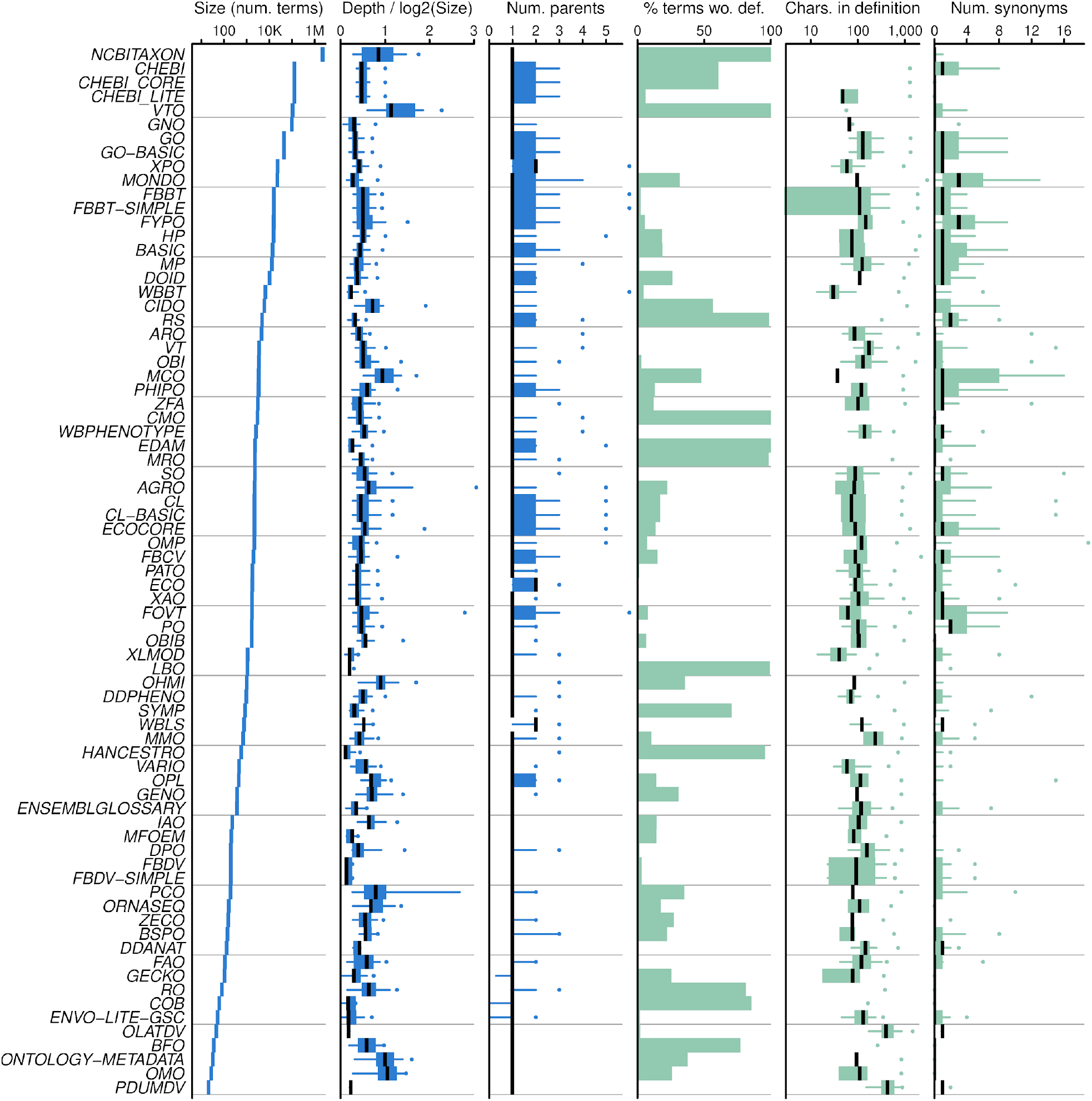
Summary statistics for selected ontologies. Vertical panels display the number of terms in each ontology, the distribution of term depths (normalized by the size of each ontology), the distribution of the number of parents for each term, the proportion of terms lacking a text definition, the distribution of number of characters in text definitions, and the distribution of the number of synonyms. Distributions are visualized by a median (central bar), interquartile range (boxes), 5%-95% quantile range (whiskers), and maximal values (dots).

In addition to structural metrics, we examined properties of text annotations attached to ontology classes. We assessed the proportion of the terms that did not contain any definition and found this spanned the full range from 0% to 100%. A hypothesis to explain missing definitions might attribute them to new terms, which may lack detail before proper curation. However, manual review suggested that missing definitions may instead be explained by maintenance goals. In taxonomies of species such as *NCBITAXON*, for example, terms systematically carried names but not any further characteristics. This suggests that detailed definitions may be considered out-of-scope for those ontologies and that terms are instead defined by their positions relative to ancestors. A related mechanism may be at play in branches of other ontologies where child nodes are very closely related to their parent. An example was the term ‘abnormal mitochondrial number’ in the human phenotype ontology, *HP*^*36*^, for which child nodes (‘increased mitochondrial number’, ‘decreased mitochondrial number’) omitted dedicated descriptions, their full meaning presumably captured by the definition of their parent.

To understand text annotations in more detail, we counted the number of characters in term definitions. These counts, which generalize word-counts and are applicable for unusual content like chemical formulae and compound words, are proxies for the information that users have to understand their meanings. Interestingly, distributions within one ontology often spanned an order of magnitude or more. However, despite the internal variability, the centers of the distributions fell in a narrow range across ontologies (25% quantile 76 characters, 75% quantile 119 characters). Thus, there is a remarkable consistency across research domains as to the amount of text that is deemed appropriate to capture unique concepts.

For a final descriptive summary, we counted the number of synonyms, i.e. alternative names, associated with ontology terms. Although synonyms are optional, the majority of ontologies employed this feature in at least some of their terms. Manual investigations of outliers revealed several approaches to their usage. In the *MONDO* ontology of human diseases, synonyms were employed to synchronize content from independent disease databases. In the plant ontology, *PO*^*37*^, several synonyms were translations between English and Spanish. In other cases such as the ontology of microbial conditions, *MCO*^*38*^, synonyms were due to terms imported from *CHEBI*, the ontology of chemical entities^39^.

Altogether, these patterns reveal that ontologies have distinctive characteristics that reflect design and curation choices. They also share similarities, not only in that they are encoded in a common file format, but also in how they use comparable amounts of text to define meaning. These similarities indicate that it is reasonable to assess the set of ontologies using a common natural-language processing and machine-learning framework.

### Ontology text annotation can train unsupervised machine-learning models

Ontologies provide vocabularies for their subject domains, definitions of relevant concepts, and relationships between concepts^1^. Although these roles are conceptually distinct, practical implementations can blur their boundaries. In particular, concept definitions can convey similarities and differences to other concepts, and thereby create an implicit graph of relationships. We set out to assess the concordance between the information encoded in the explicit ontology hierarchies and the implicit graphs created by text-based definitions. Some approaches have compared lexical and semantic annotations in case studies^25–27^. Here, we aimed to perform an analysis at the pan-ontology level. Given the amount of text associated with ontology terms was stable across many ontologies, we reasoned that these definitions could be the basis for a systematic comparison.

For our calculations, we encoded ontology terms into numeric vectors through k-merization. Such straightforward encodings are known to perform well for classification tasks - only marginally worse than deep neural networks^40^ - and can be used on both small and large datasets. We used software that has already been demonstrated to yield reasonable mappings across two ontologies^41^. It is thus suited to process ontology data, both in terms of capturing rich concepts and in terms of computational efficiency. However, other approaches to encode text and ontology terms are possible^14,15,17^.

We generated three datasets for each ontology (Figure 3A). The simplest version associated each ontology identifier with the term’s name, only. A second version comprised the term name, definition, synonyms, and comments. This attempted to collect a comprehensive summary of each ontology term, as provided by the project curators. In a third version, the descriptions were further supplemented with the names and definitions of parent terms. This version thus incorporated connections between terms that would normally be encoded via the ontology hierarchy. The three data versions were denoted ‘names’, ‘full-text’ (FT), and ‘full-text plus parents’ (FT+), respectively.

**Figure 3.**
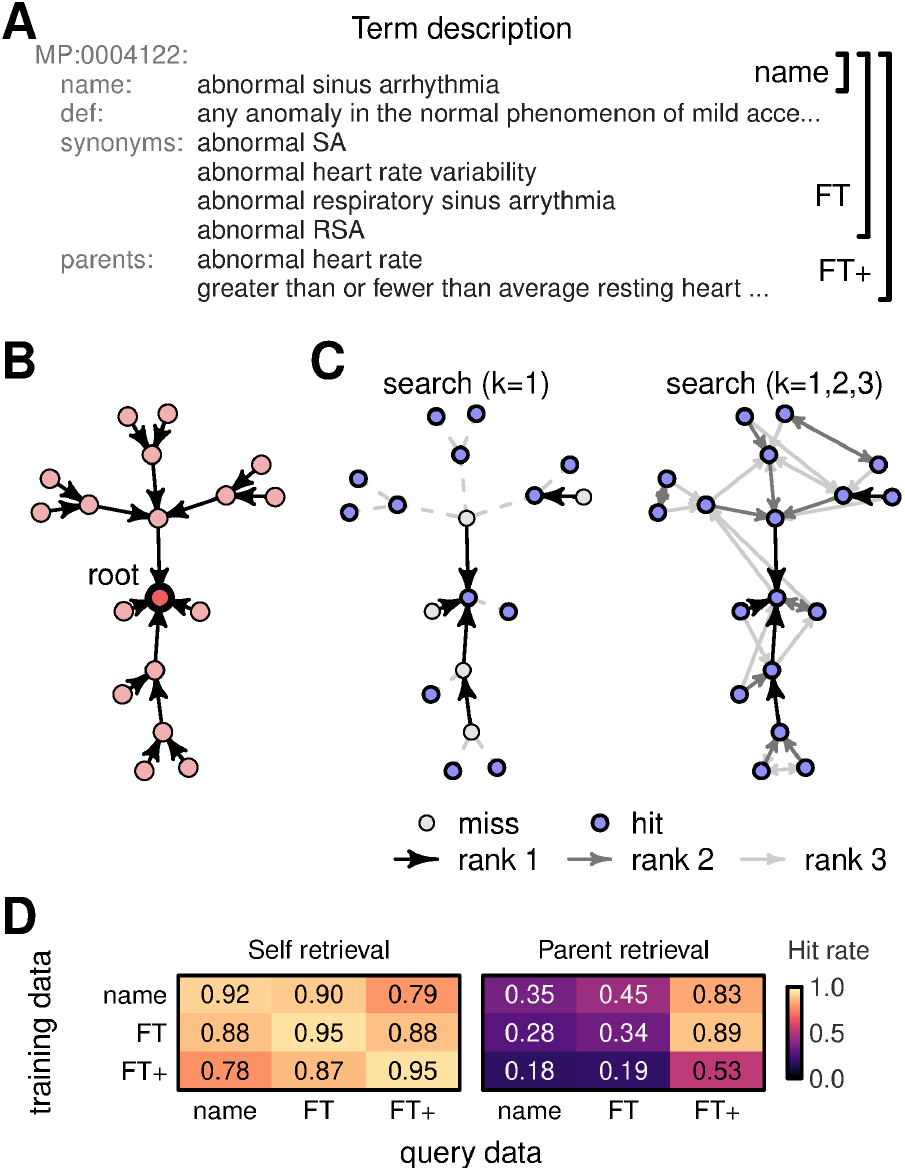
Reconstruction of ontologies hierarchy. (A) Example of text associated with an ontology term from the mammalian phenotype (MP) ontology. One version includes only the term name, another version includes full-text (FT), and another has full text plus definitions of parent terms (FP+). (B) Graph for a branch of the MP ontology. Dots represent ontology terms, arrows represent directional ‘is a’ relationships. The dark dot is the root node. (C) Example of an ontology-reconstruction calculation using nearest-neighbor search. The first view shows first-ranked predictions (k=1); the second view shows the top three predictions (k=1,2,3). Labels ‘hit’ and ‘miss’ indicate when text from a given node is matched to itself via search. (D) Summary of reconstruction performance averaged over the selected ontologies.

For each ontology, we parsed the FT dataset into k-mers along with an auxiliary dataset based on an English dictionary (www.wiktionary.org). By counting k-mer frequencies in such mixed data, we produced weights that capture usage in common language but also include domain-specific nuances. Next, three unsupervised machine learning models were constructed for each ontology using the name, FT, and FT+ datasets. Training consisted of parsing the datasets into k-mers, encoding data into numeric vectors, and generating nearest-neighbor (NN) indices^42,43^. We then used data from the same three datasets as queries for NN searches and compared outputs to expected mappings. These calculations represent a (quasi-)circular design whereby the same data is used for model training and evaluation. Therefore, they cannot be used to assess the quality of the models according to common metrics. However, search outputs can reveal informative patterns about the relatedness between ontology terms. This bears similarity to evaluations of unsupervised clusterings.

To illustrate our approach, we performed the procedure on a branch of the mammalian phenotype ontology, MP^44^, pertaining to ‘abnormal heart rate’ (Figure 3B). The branch consisted of 18 terms, including ‘abnormal sinus arrhythmia’ (ASA) as a child term to the root. We created three datasets - name, FT, and FT+ - with 18 terms each and trained three separate models. In a search using the ASA name against a model trailed with name data, we expected the model to return a mapping to that term’s identifier. Indeed, this was the case empirically, and analogous outcomes were also repeated for the other 17 terms in the small example. These results indicate that our approach was at least 95% accurate despite using approximate (as opposed to exact) NN search. Next, we queried the model trained on FT data with text from the FT+ dataset. Four items were mapped at k=1 to other identifiers, and extending the search to k=2,3 nearest neighbors revealed a network of interconnections (Figure 3C). The interpretation of the empirical, machine-generated graph is subtle but informative. On the one hand, searches using FT+ data were primed to produce mappings to ontology terms and their immediate parents. Thus it is not surprising that the k=1,2,3 graph resembled the ontology hierarchy. On the other hand, some empirical search results deviated from the expected pattern, in some cases mapping to sibling terms or even further relations, indicating possible sources of tension.

To study such properties in a systematic fashion, we assessed outputs of NN searches on the entire set of ontologies, considering up to five neighbors (k=1, 2, … 5). We summarized results by measuring the proportions of terms that could be mapped to their respective objects (self-retrieval) and their parents (parent-retrieval) using the nearest neighbors. We assessed these measures at the ontology level and then averaged over the set of ontologies (Figure 3D). Averaged performance for self-retrieval was 93-96% when the training and query datasets were the same. Performance dropped when the training and query data were technically distinct. However, it remained above 75% even when models were trained with FT+ data but were provided only with term names during querying. This indicates stability to perturbations.

Performance on the parent retrieval task was considerably lower (Figure 3D). Using FT data for training and querying, i.e. not including any text from parents, the averaged score was 34%. This serves as a baseline for the capability of a simple model to capture the graph structure from text definitions alone. The best average parent-retrieval, 89%, was achieved by training with FT data and querying with FT+ data. This calculation was primed to perform well on the parent-retrieval task - it provided the model with text describing the expected term-parent relationships and the performance assessment considered up to five nearest neighbors. Despite this, some text definitions created confusion for the machine models. Low parent-retrieval performance for name-based queries further suggests that the machine-generated representations create connections between terms that are distinct from ontology hierarchies.

### Ontologies have informative definitions but incomplete sets of relations

To gain further insight into the performance on the self-retrieval and parent-retrieval tasks, we investigated scenarios using FT datasets at the ontology level. We explored correlations between performance metrics and ontology characteristics (Figure 4). Modeling with multiple regression revealed several variables around the nominal level for statistical significance. However, due to the small dataset size, multiple hypothesis testing, and changing importance levels in different modeling approaches, this could not provide clear-cut insights. Moreover, visualizations of the data revealed that mild trends were overshadowed by outliers.

**Figure 4.**
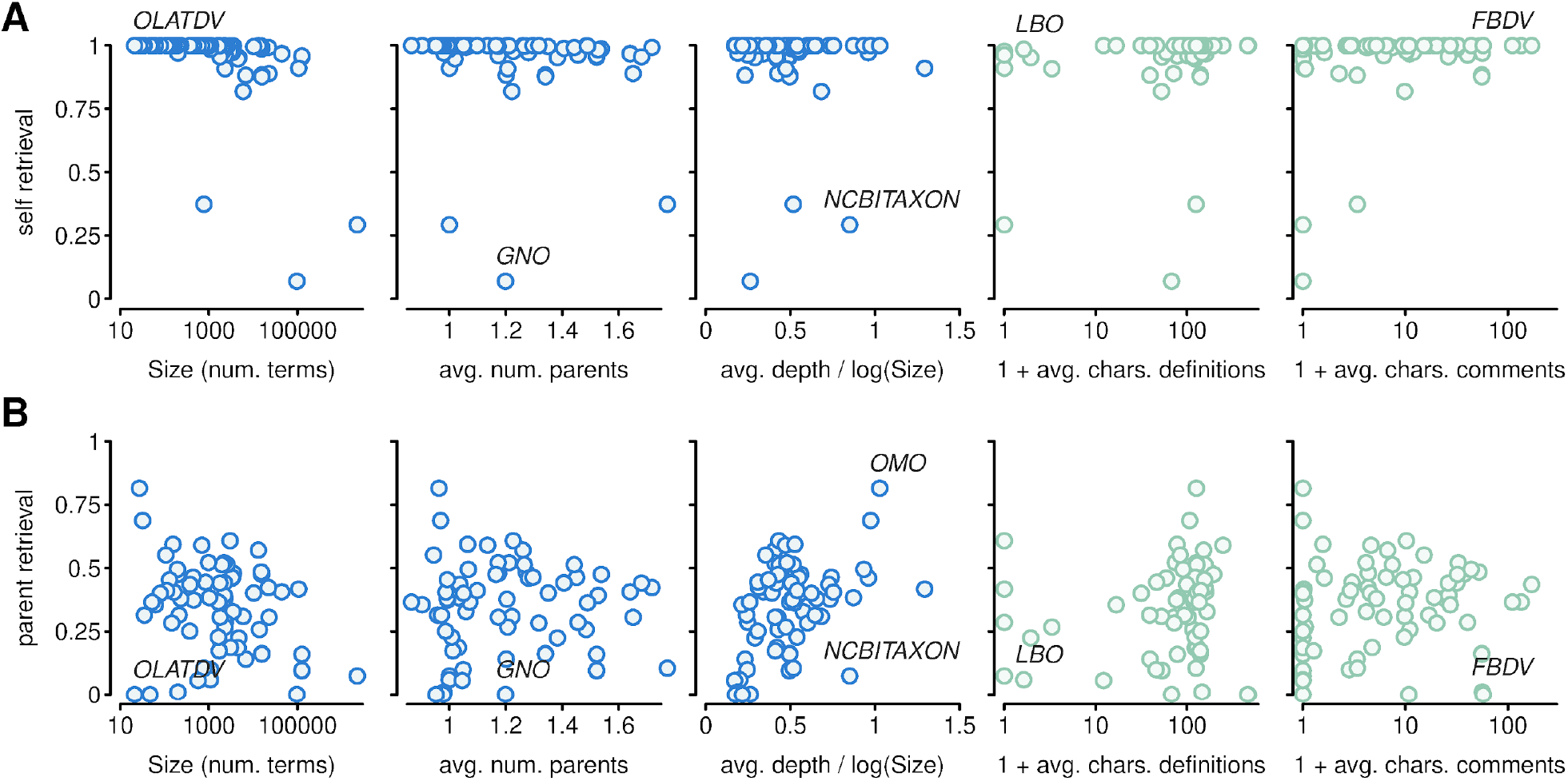
Factors affecting ontology reconstruction from text descriptions. Panels show relations between scores from machine learning models, trained and queried with FT data, and various explanatory variables. Vertical axes show (A) self-retrieval and (B) parent-retrieval scores. Dots represent ontologies. Selected dots are labeled manually.

We first inspected the ability of models to map term descriptions to their source terms. The distribution of outcomes was narrow (mean 0.96, IQR 0.97-1.00), but there were a small number of outliers (Figure 4A). The lowest performance was recorded for *GNO*, an ontology of glycan names. Despite this ontology recording definitions for most of its terms, manual inspection revealed that many definitions held very similar k-mer profiles with variations only in numeric literals. Another poor performer was *NCBITAXON*, the taxonomy of species, which systematically omits term definitions. The limited number of k-mers associated with each term may have limited performance in NN searches. However, low annotation levels alone were not predictive of poor performance and, for example, *LBO* (livestock breeding) achieved a high score despite limited text annotations. Altogether, these observations suggest that strong outliers may be improved through increasing the sensitivity of nearest-neighbor search, but that otherwise our modeling approach adequately captured the meaning for most ontology terms.

Parent-retrieval performance was more dispersed (mean 0.34, IQR 0.21-0.46) and revealed a different set of outliers (Figure 4B). The best-performing ontology was *OMO*, a project intended to understand ontology development itself. *OMO* combined a small size, well-annotated terms, and a deep hierarchy. However, those properties were not sufficient individually to give other ontologies similar performance. Indeed, the lowest nominal outcome was for *OLATDV*, an ontology of the life stages for Medaka, which is of similar size to *OMO*. Another similar case was *FBDV*, an ontology of Drosophila development. Low parent-retrieval outcomes for *OLATDV* and *FBDV*, which had very good self-retrieval, were caused by their shallow depths. Terms in those ontologies were more often mapped to siblings than to the parents: specific life stages were mapped to neighboring life stages rather than to an abstract parent. Other low-scoring ontologies included *GNO* and *NCBITAXON*, again likely due to limited term descriptions.

In summary, results from the self- and parent-retrieval tasks validated that ontology term definitions provide distinctive content that can be learnt by machine learning models. Self-retrieval was consistently high and aggregate averages were only driven downward by a small number of peculiar outliers. Parent-retrieval was higher than expected by chance - the expected random performance is on the order of the inverse of the ontology size - but it was nonetheless far from perfect. To the extent that low scores may be due to crude k-mer representations, these results can be treated as baseline levels for comparison for more sophisticated approaches. However, given high self-retrieval performance even with perturbed data and variability in the parent-retrieval, these results suggest that ML models learn distinct relations between terms than those provided by the ontology hierarchies. In turn, this implies that the formal ontology hierarchies may be incomplete, or term definitions confusing. Irrespective of whether these differences arise because of artefacts during ML training or due to difficulties in capturing complex biological concepts in ontology hierarchies, there are opportunities to use machine-learning models to feed back to ontology development. Details of such feedback must be domain-dependent, so we next performed case studies.

### Machine learning models suggest ontology modifications

As a first example for how machine learning may suggest ontology modifications, we considered the ontology for Drosophila development, *FBDV*^*45*^, which was an outlier in our parent-retrieval calculations. The ontology is a relatively small set of 200 terms with a simple hierarchy and non-empty term definitions (c.f. Figure 2). But its terms are organized in a shallow structure and our nearest-neighbor searches often mapped terms to siblings rather than to parents. To gain further insight, we used the outcomes of our search calculations to create a UMAP visualization in a two-dimensional space^46^. This approach is similar to studies of node embeddings^15–17^ in that it projects similarities between nodes into a low dimensional space. The visualization revealed that terms cluster into more groups than there are branches in the ontology (Figure 5A). This observation was stable to changes in embedding hyperparameters, but did not apply for embeddings constructed from the ontology graph alone using a graph layout algorithm, node2vec^15^, or UMAP based on graph distance (Supplementary Figure S3). Upon manual inspection, clusters corresponded to embryonic, larval, pharate, pupal, and other stages of fly development. They thus identified biologically relevant groups despite these not being formally separated via ‘is a’ relationships in the ontology definition. The new clusters can be seen as suggestions to create intermediate nodes. Thus, machine-driven analyses of text annotations provide actionable insights for ontology maintenance.

**Figure 5.**
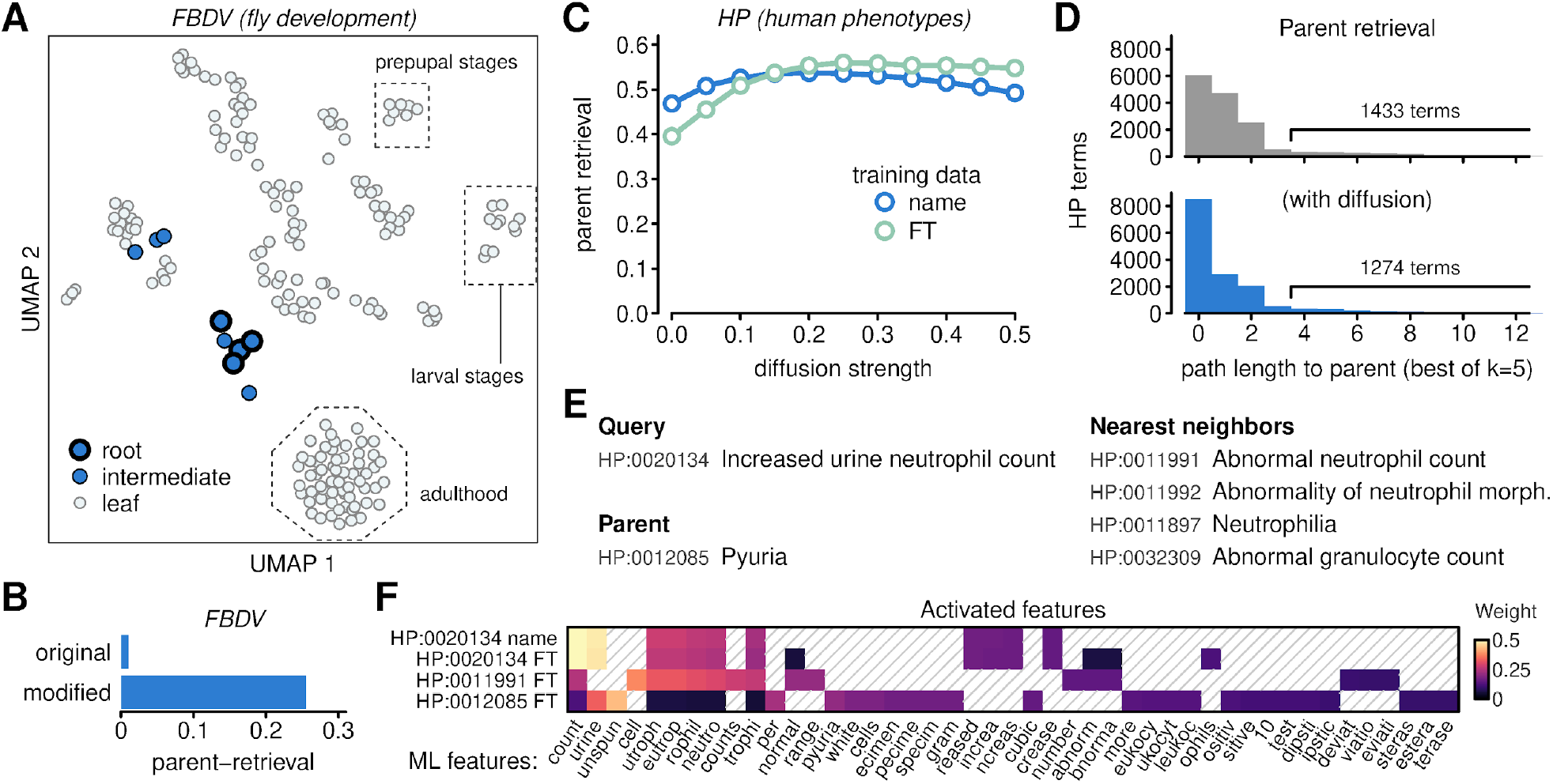
Feedback from ML for ontology development. (A) Unsupervised UMAP embedding of *FBDV* ontology terms (dots) according to their similarity of text-based annotations. Selected groups of terms are labeled manually. (B) Parent-retrieval performance in the original FBDV ontology and a modified version with more hierarchical relations. (C) Effect of diffusion on parent retrieval in the human phenotype ontology, *HP*. Dots represent measurements and lines are interpolations. Two series represent results on models trained with names and FT datasets; all queries performed with names data. (D) Summary of errors in parent retrieval without (above) and with (below) diffusion. (E) Example of an ontology term with a large parent-retrieval error. (F) Activation heatmap for exemplar term representations.

Interestingly, some of these suggested patterns appear to already be captured by the use of ‘substage of’ relationships in the *FBDV* ontology. To test the extent this additional hierarchy may impact on our parent-retrieval scores, we created a modified version of the ontology by replacing ‘substage of’ relationships by ‘is a’ relationships so that they may be included in our calculations. This is equivalent to extending data-parsing and evaluation procedures to treat all ontology relationships with equal weighting. We observed that parent-retrieval performance indeed increased - from near zero to 26% (Figure 5B). A similar modification in the context of the gene ontology, *GO*^*3*^, using ‘part of’ relationships on a par with ‘is a’ links, increased apparent parent-retrieval performance from 41% to 45%. These changes illustrate that evaluations can be sensitive to how ontology definitions are parsed. But because parent-retrieval performance for *FBDV* remained far from perfect, patterns captured by the machine learning model remained different from those explicitly encoded in the ontology.

As another case study, we analyzed the *HP* ontology of human phenotypes^36^. This ontology was larger (15,054 terms) and organized in a deeper structure (median depth 7, maximal depth 14). It performed well in our calculations for self-retrieval and in-line with other ontologies for parent-retrieval. We reasoned that it might be possible to improve parent-retrieval by imputing features. An implementation based on diffusion was previously described to improve mappings across two ontologies^41^. To assess the impact of this approach for parent retrieval, we repeated the search calculations using only term names as inputs, applied imputation based on FT+ annotations, and then queried against models trained with name and FT data. Diffusion effectively augmented queries with additional text on the basis that the new features co-occurred in term-parent relationships. Thus, it steered the search process to associate terms to their ontology parents. We repeated this procedure using various diffusion strengths and indeed observed an improvement in parent-retrieval (Figure 5C), although at the cost of reduced self-retrieval (Supplementary Figure S4). The most pronounced effect occurred when querying against models trained with FT annotations: parent-retrieval increased from 40% to 56%, and this improvement was stable for a wide range of diffusion strengths.

Despite better parent-retrieval after imputation, many *HP* terms were still not mapped to their parents. We investigated further by computing the distances - in terms of the ontology graph - between our search results and the parent terms that we aimed to retrieve (Figure 5D). This confirmed that imputation reduced path lengths, and further showed that more than 90% of *HP* terms were mapped to terms within three graph steps from the parent. Thus, even when searches did not return the parent term, they returned terms in related branches.

Interestingly, distributions of path lengths contained long tails of *HP* terms that were mapped far away from the parent (Figure 5D). We inspected individual terms that gave high path-length errors in both undiffused and diffused settings. One example was the term ‘increased urine neutrophil count’ for which the ontology parent is ‘Pyuria’ (Figure 5E). Because those names do not share any substrings, it is unsurprising that a search against ‘name’ annotations did not return the parent term. However, search against the FT corpus did not return the parent term among the top five results either, even though the parent does include synonyms that link pyuria to neutrophils in the urine. Instead, the leading search result was ‘abnormal neutrophil count’ with a path-length error of 9 (Figure 5E). While this hit appears to be appropriate, its definition restricts its meaning to the context of the circulatory system and it is set in a different branch of the ontology. Other hits reported by the NN search were also blood-related phenotypes.

To gain insight into how the parent-retrieval error might be resolved, we visualized the activated features (k-mers) associated with the query term, expected parent, and the leading search result (Figure 5F). This revealed the importance of individual parts of the query in the match against the database. While k-mers making up the word ‘neutrophils’ were present in the definition of ‘Pyuria’, their contribution was masked by other k-mers originating from a description of the experimental procedure that is used to detect the phenotype (c.f. k-mers associated with ‘urine specimen’, ‘Gram staining’, ‘dipstick test’). The drill-down highlights the aspects of the definition that is picked up by automated tools and this knowledge can be used in two distinct ways. It might be used to adjust the machine learning models, for example through user-driven learning^41^. Alternatively, the insight can be incorporated into the ontology itself, for example by editing term definitions in order to reduce the chance of misleading ML models. These are practical mechanisms for ontology development, and similar investigations can be repeated for other poorly-performing terms (Supplementary Figure S5).

Finally, we looked at ontology nodes near the leading search result ‘Abnormal neutrophil count’. While this term is not an ideal match for our query due to the distinction between blood and urine, its parent is ‘Abnormality of neutrophils’ and is agnostic of the biological context. The more general term is thus, arguably, an appropriate match for ‘increased urine neutrophil count’. Had the ontology included a link between ‘Pyuria’ and ‘Abnormality of neutrophils’, this example would have been evaluated with a path-length error score of 2 instead of 9. The nonzero score would have still conveyed an imperfection in the match, but not of a magnitude to suggest unrelated phenotypes. ML search may thus be a mechanism to identify branches in the ontology that should be linked closer together - in this case, bridging homeostasis and the immune system. The technique of linking terms to more than one parent is already widely used to capture complex concepts (c.f. Figure 2), so this ML-driven adjustment is feasible in practice.

## Discussion

Ontologies help to organize knowledge in research areas as diverse as chemistry and ecology. They also adapt in response to the needs of their communities. This work addressed the possibilities for knowledge transfer between ontologies and machine-learning models, in particular from the perspective of dissecting ML models in order to streamline ontology development. Our approach included a survey of the ontology landscape and highlighted practical, actionable strategies that are applicable across all ontologies.

Despite common data repositories and file formats, there is heterogeneity in the practical architectures of ontology projects. To some extent, this reflects the breadth of topics, as well as the disparate needs and priorities of the respective research communities. However, heterogeneity can also impact on how well insights from one application can translate to other contexts. In particular, two distinct aspects can have an impact on machine learning.

One aspect of heterogeneity lies in the usage of relationships between ontology terms. Hierarchical ‘is a’ relationships are ubiquitous in ontology visualizations, and they serve to propagate information from specific to broader terms in applications. For example, they are exploited in calculations performing ontology matching and scoring genotype-disease associations^47^. Some hierarchy-driven classifiers outperform all other approaches in situations with scarce training data^19^. Our pan-domain review highlights that ontologies range from very shallow to very deep, so this heterogeneity will likely limit the ability to transfer ML modeling techniques tested on deep ontologies to shallow domains. Interestingly, a case study with the *FBDV* ontology illustrates that embeddings based on text annotations can learn rich and informative patterns beyond what is encoded in explicit hierarchical relations. Indeed, embeddings can suggest ways to increase the depth of an ontology through subcategories. The same case study also emphasizes that when there are multiple relationship types, i.e. relations other than ‘is a’ hierarchies, the way this information is passed to ML models can have a substantial impact on performance metrics. Ontologies that utilize nontrivial relations may consider including recommendations for how to use these within ML models.

Another aspect of heterogeneity pertains to differences among terms within individual ontologies. In this regard, we generalized previous explorations of semantic-lexical consistency^25–28^. We performed a series of calculations in which we used term names to query ML models to identify parents. Our models returned the correct parent terms among the top five outputs for fewer than half of the inputs. This indicates there are discrepancies in the information encoded via text definitions and graph-based relations. In a case study using the *HP* ontology, we could increase retrieval from 40% to 56% by using automated feature imputation, and to around 90% by accepting fuzzy matches. The fact that performance was reasonable for a majority of terms but remained poor for a considerable fraction suggests that the discrepancies are not entirely artifacts of our modeling approach. Rather, the calculations pinpoint areas that are challenging for machine understanding. The same characteristics may create bottlenecks in other applications, so identifying and avoiding them would increase the value of the ontology in downstream applications.

Dissecting machine-generated representations of ontology concepts can lead to adjustments through two distinct mechanisms: modification of text annotations and addition of relationships. We showed that activation maps of k-mers weighted by their information-content can be informative and interpretable. This technique illustrated, for example, that a description of ‘Pyuria’ in the human phenotype ontology *HP* is dominated by k-mers unrelated to biological characteristics. This information is a prompt to modify the definition in order to place more emphasis on the molecular and cellular characteristics (as is the convention in the *HP* ontology). Importantly, the computational cost of such alerts is minimal. A more involved diagnostic of internal heterogeneity relies on parent-retrieval. The idea behind this technique is that machine-driven representations should be capable of capturing hierarchies. In the case of the ‘Pyuria’ phenotype, this reasoning suggested forming a new link between two otherwise distinct ontology branches (blood and urine phenotypes). Such a connection could not be inferred from logical reasoning based on ontology axioms alone.

The idea of adjusting ontologies based on ML-based analyses holds both promise and potential pitfalls. A key advantage is simpler training of new ML models for downstream applications: when information in ontology hierarchies and ML-derived representations are consistent, then biological datasets can be utilized to uncover biological patterns rather than to correct ontological artifacts. Given that biological datasets can be limited, this factor can be important in practice. Potential pitfalls of adjusting ontologies based on ML feedback include “over-fitting” definitions to one particular algorithm, or even sacrificing biological nuances to achieve better performance in a benchmarking test. Any adjustments must therefore rely on judicious curation.

In summary, we argued that as more biological pipelines employ ML components, it is worthwhile to dissect how these techniques utilize ontology annotations. We showed that ML techniques can create specific insights that can be incorporated into ontology definitions through editing text annotations and adding new relations between terms. Both mechanisms are compatible with current curation practices and are thus feasible in practice. Such feedback can create a reinforcement cycle through which ML models better utilize ontology data during training and thus deliver superior performance in downstream applications.

## Methods

### Pan-ontology dataset

Ontology files were downloaded from the OBO foundry (http://www.obofoundry.org) and the OLS (https://www.ebi.ac.uk/ols). Ontology definitions were downloaded in the obo file format, when available. This is a common baseline format for many ontologies. For ontologies that were available only in owl format, the owl files were downloaded and converted to obo using the pronto library (https://github.com/althonos/pronto). Ontologies that failed owl-to-obo conversion, or were only available in other file formats, were omitted from further processing.

### Machine-learning

Machine-learning models for individual ontologies were built using crossmap (https://github.com/tkonopka/crossmap), a framework for analyzing text-based data^41^. As preparation, obo files were parsed to extract term names, definitions, synonyms, and ‘is a’ relationships. Ontology terms marked as obsolete were omitted from analysis. The parsed data were arranged into three distinct collections, denoted ‘names’, ‘FT’, and ‘FT+’. The ‘names’ collection associated each ontology term with its name, only. The ‘FT’ associated terms to their full-text description, including name, definition, synonyms, comments, and the name of the top-level term. The third collection, ‘FT+’, extended the full-text description with the name and definition of parent terms.

For each ontology, the ‘FT’ data collection was first used together with a collection from the wiktionary (www.wiktionary.org) to estimate the frequency of k-mers (k=6). These frequencies were then used to build a crossmap knowledge-base holding the ‘names’, ‘FT’, and ‘FT+’ collections. The two-pass approach is a means to weight k-mers in a domain-specific manner.

The crossmap knowledge-bases were queried using the batch search feature. Search was carried out requesting 5 hits; all other settings were left at their defaults. Outputs of searches from crossmap knowledge-bases were analyzed in R using custom scripts.

## Data availability

Scripts are available at https://github.com/tkonopka/pan-ontology-ml. A dataset with raw ontology definitions, processed data used in the analysis, and outputs from calculations is available at zenodo.org (10.5281/zenodo.4311237).

## Acknowledgements

We would like to thank Chris Mungall for comments on a manuscript draft.

## Supplementary figures

**Figure S1.**
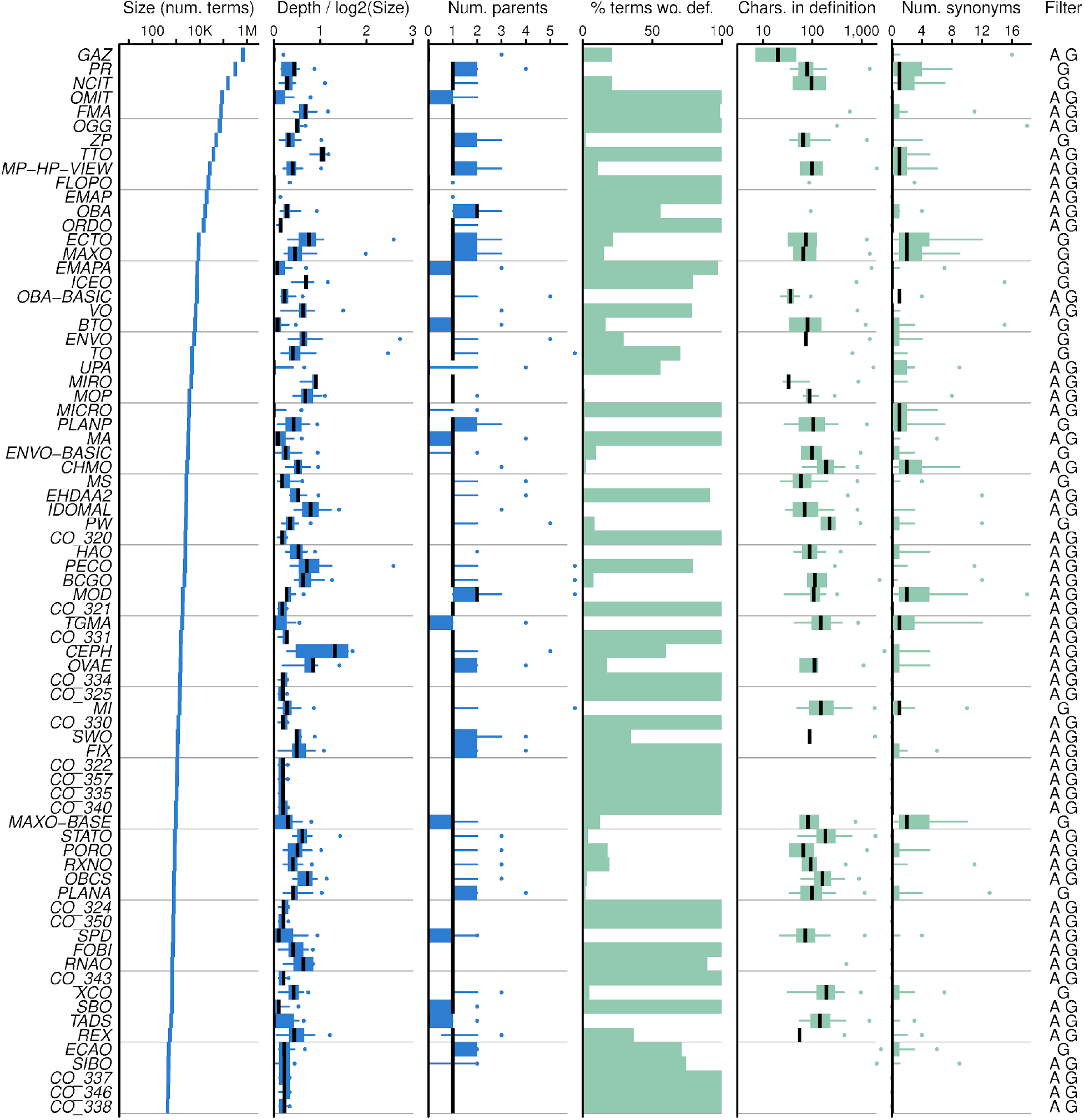
Summary of additional ontologies available through the OBO foundry and OLS, but not selected for in-depth analysis (1/2). Ontologies are summarized by the total number of terms, the distribution of term depths (normalized by the size of each ontology), the distribution of the number of parents for each term, the proportion of terms lacking a text definition, the distribution of number of characters in text definitions, and the distribution of the number of synonyms. Distributions are visualized by a median (central bar), interquartile range (boxes), 5%-95% quantile range (whiskers), and maximal values (dots). The filter code indicates criteria for omitting the ontologies from in-depth analysis: non-recent age (A), graph structure (G), small size (S).

**Figure S2.**
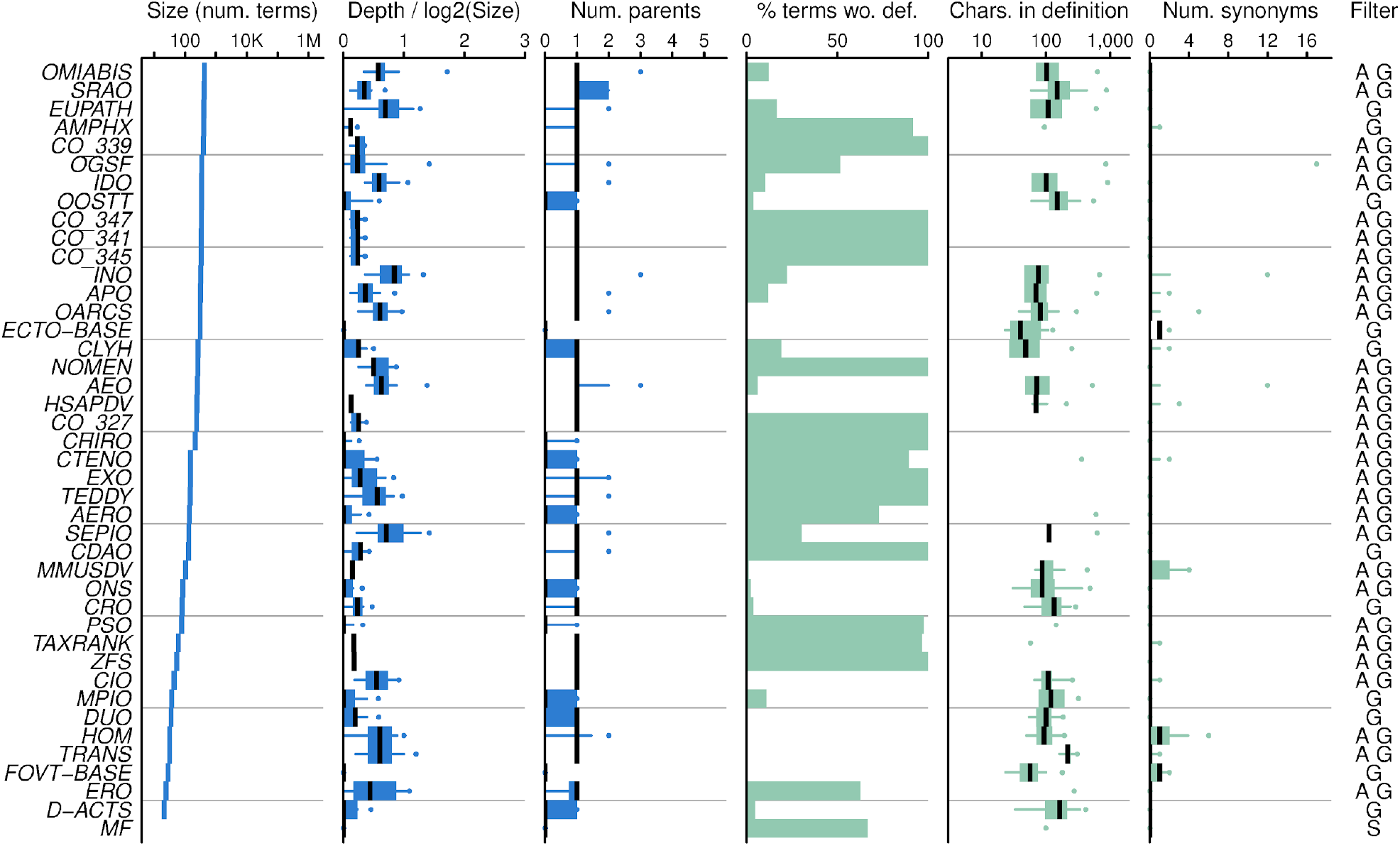
Summary of additional ontologies available through the OBO foundry and OLS, but not selected for in-depth analysis (2/2). Continued from previous figure.

**Figure S3.**
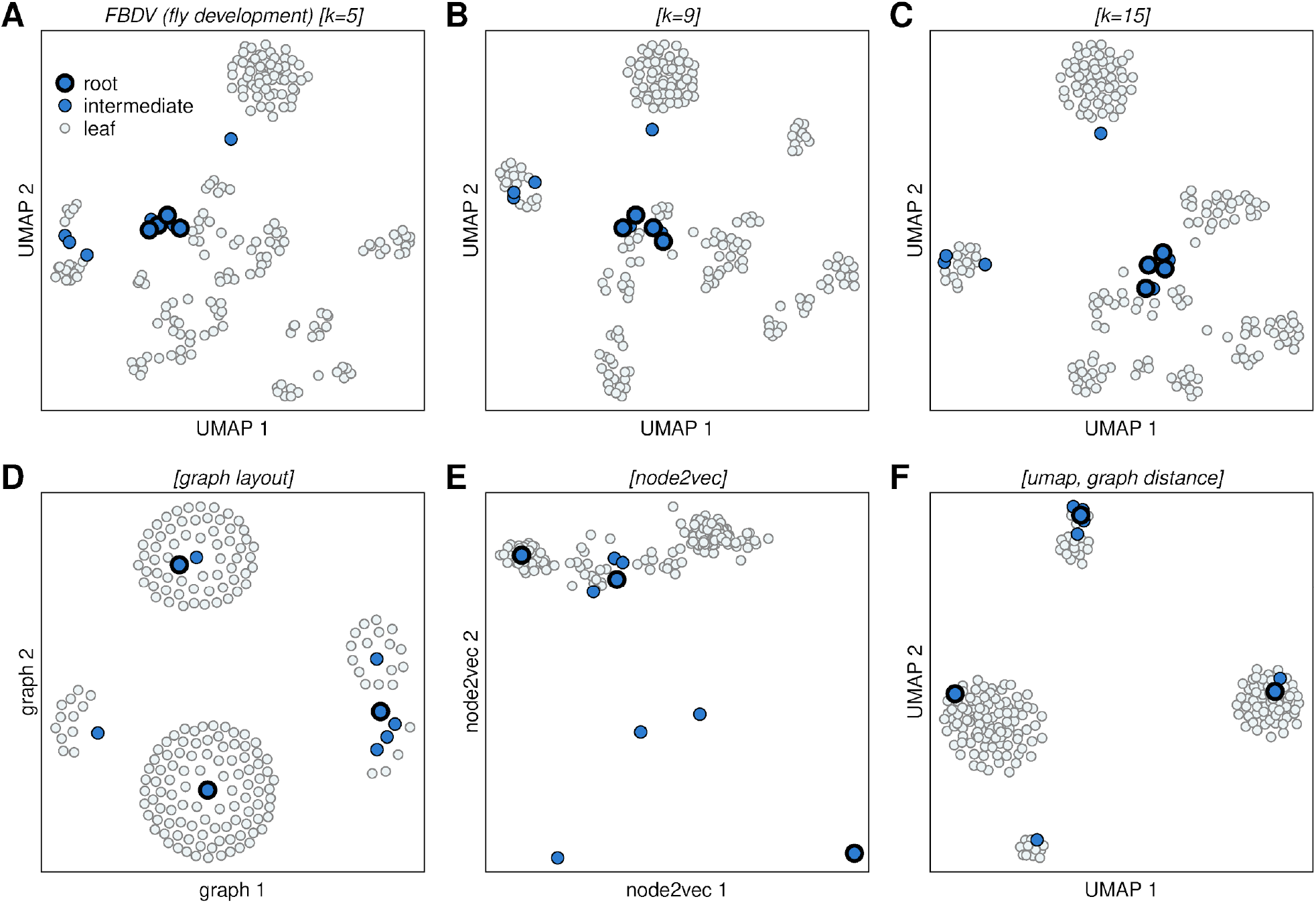
Embeddings of FBDV ontology terms. All panels represent embeddings of FBDV ontology terms into an abstract 2D space. Dots represent ontology terms and are colored by their position in the ontology hierarchy as roots, intermediate nodes, and leaf nodes. (A-C) UMAP embeddings constructed from semantic distances based on text-annotations. Panels differ in that neighborhoods are computed using (A) k=5, (B) k=9, and (C) k=15 neighbors. The embedding with k=5 is conceptually equivalent to the embedding in one of the primary figures, but differs because of stochasticity. (D-F) Embeddings constructed from distances based on the ontology graph structure. Panels generated using (D) a graph layout algorithm, (E) node2vec with default arguments, and (F) UMAP based on graph distance with k=15.

**Figure S4.**
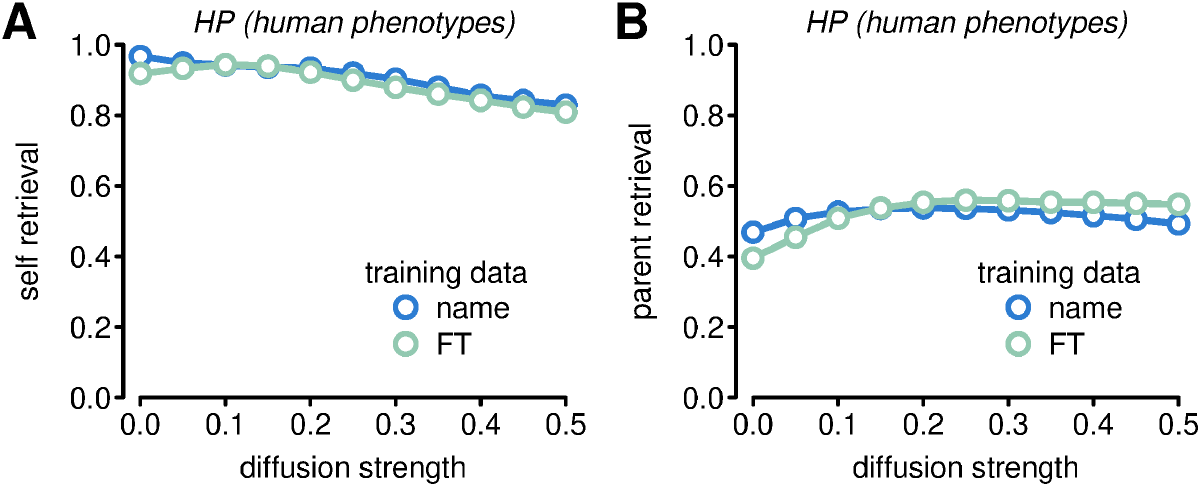
Effect of diffusion on self-retrieval and parent-retrieval performance in the HP ontology. Panels summarize calculations that use HP ontology term names as queries against models trained with HP full-text (FT) data. Diffusion is driven by HP FT+ data. Panels show effects on (A) self-retrieval and (B) parent-retrieval. Panel (B) reproduces one of the panels in the primary figures.

**Figure S5.**
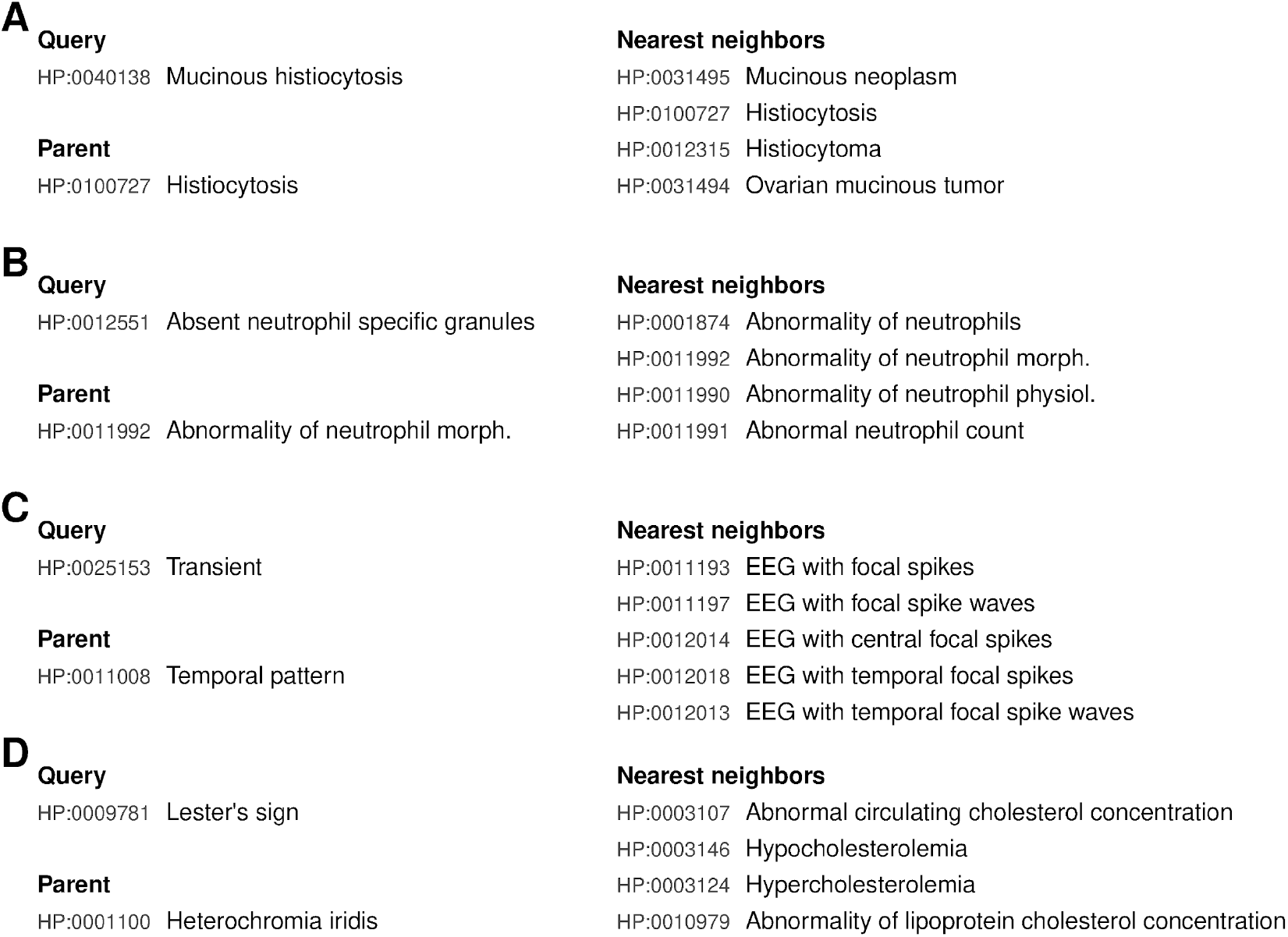
Examples of HP term mappings that are substantially different from the expected location in the ontology. Panels (A-D) show individual query terms, parent terms defined by the ontology, and outcomes of nearest neighbors searches. All searches were performed using FT+ diffusion.

